# OryzaG3: A Single-species Genomic Foundation Model Pretrained on Rice Pangenome

**DOI:** 10.64898/2026.05.22.727045

**Authors:** Lin Yang, Yu Xia, Zhuang Yang, Chengcai Xia, Tingxun Wu, Meiling Zou, Zhiqiang Xia

## Abstract

While multi-species genomic language models have advanced biological representation learning, high-quality, single-species foundation models for crops remain scarce. Leveraging recently expanded rice pangenome resources, we introduce OryzaG3, a species-specific DNA language model with 700M parameters. OryzaG3 was pretrained on 59.20 Gb of chromosome-level sequences from 149 high-quality rice genomes using a non-overlapping 3-mer tokenization strategy and a causal language modeling objective, featuring context-length variants up to 32k tokens. On the Plants Genomic Benchmark polyA prediction task, OryzaG3 achieves competitive predictive performance against leading multi-species models while delivering a four-fold increase in inference throughput under identical long-context conditions. Ultimately, OryzaG3 demonstrates that lightweight, single-species foundation models trained on high-quality pangenomes can match multi-species benchmarks while significantly reducing computational overhead. This work provides a scalable framework for rice functional genomics, molecular breeding, and targeted crop foundation model development.

## Introduction

Genomic language models (GLMs) have recently revolutionized biological representation learning by capturing complex features directly from massive DNA sequences through self-supervised pretraining^1^. These studies have achieved breakthroughs in downstream tasks such as gene structure prediction, regulatory element identification, and variant effect assessment, demonstrating that motifs, local syntactic structures, and evolutionary constraints can be effectively mastered at scale^2–10^.Currently, research in this field has primarily focused on animal and human genomes, with representative works including DNABERT^3^, Nucleotide Transformer^4^, and EVO^5^.

In contrast, plant genomic pretrained models are still in their early stages of development. Existing plant genomic pretrained models, such as AgroNT^11^, PDLLMs^12^, and Botanic0^13^, largely focus on multi-species mixed training, while dedicated single-species models remain scarce^14^. These models can learn relatively general plant genomic representations by leveraging cross-species sequence information^11–13^, but their ability to capture intra-species variation, local selection signals, and species-specific regulatory sequences in particular crops may still be limited. Moreover, large-scale multi-species models typically have high parameter counts and inference costs, posing efficiency challenges for practical deployment in crop breeding and species-specific tasks.

Rice (*Oryza sativa*), one of the world’s most important staple crops and a model species in plant genomics, has accumulated rich high-quality multi-omics data^15^. Recent high-quality rice pangenome studies have released chromosome-level genomes for 149 cultivated and wild rice accessions, encompassing rich genetic diversity from wild relatives to modern varieties^16^. This provides an ideal data foundation for developing single-species dedicated models. Concurrently, model architectures in natural language processing are evolving toward lightweight and high-efficiency designs^17^. Open-source model architectures have achieved strong modeling capabilities at relatively small parameter scales^18^, with efficient Transformer^19^ architectures, Rotary Position Embeddings (RoPE)^20^, Grouped Query Attention (GQA)^21^, and other designs offering valuable technical foundations for long-sequence DNA modeling.

Here, we present OryzaG3, a single-species DNA language model pretrained on a comprehensive pangenome comprising 149 high-quality rice accessions. Adopting a non-overlapping 3-mer tokenization strategy paired with a causal language modeling (CLM) objective, we developed two model variants featuring context lengths of 8k and 32k tokens (OryzaG3-8k and OryzaG3-32k). We systematically evaluate OryzaG3 on the rice polyA prediction task within the Plants Genomic Benchmark (PGB)^11^, demonstrating that it achieves downstream transferability comparable or locally superior to leading multi-species plant models, such as AgroNT^11^, Botanic0-L^13^, and OneGenome-Rice^14^, while dramatically improving inference throughput. Furthermore, by tracking downstream performance across various pretraining stages, we reveal how pretraining sufficiency dynamically shapes functional genomic representations. Finally, we provide empirical benchmarks on the computational velocity and memory trade-offs enabled by FlashAttention-2^22^ and gradient checkpointing^23^, while discussing the scalability of RoPE and hybrid attention mechanisms for ultra-long-context genomic modeling. Together, this work establishes a resource-efficient, transferable technical paradigm for constructing lightweight, species-specific foundation models across diverse agronomic crops.

## Results

### Construction of the single-species rice foundation model OryzaG3

We developed OryzaG3, a 700M-parameter DNA language model, to learn sequential representations of the rice genome. The pretraining corpus derives from 149 chromosome-level genomes in the latest high-quality rice pangenome study, covering cultivated rice, wild rice, and related outgroup species^16^. Only sequences from chromosomes 1-12 were extracted. After data standardization (including cleaning, N-base handling, and filtering), the final training corpus totaled 59.20 Gb. This corpus deeply reflects the intra-species and inter-species genetic diversity within the Oryza genus.

To accommodate DNA sequence characteristics, OryzaG3 employs a non-overlapping 3-mer tokenization strategy, converting genome sequences into token sequences with a 3-bp stride. The final vocabulary size is 96, comprising 64 standard 3-mer tokens and several special tokens. Compared with the natural language vocabulary, OryzaG3’s compact rice genome-specific vocabulary significantly reduces embedding layer parameters and improves the compactness of sequence encoding.

OryzaG3 uses a CLM^24^ objective for self-supervised pretraining, predicting the next token based on observed upstream 3-mer tokens. We trained OryzaG3-8k and OryzaG3-32k from scratch to meet the analytical needs of both standard-length and ultra-long DNA sequences. The overall model construction and evaluation workflow (including pangenome sequence preprocessing, tokenization, self-supervised pretraining, checkpoint evaluation, downstream fine-tuning, and computational efficiency analysis) is shown in Fig. 1.

**Fig. 1.**
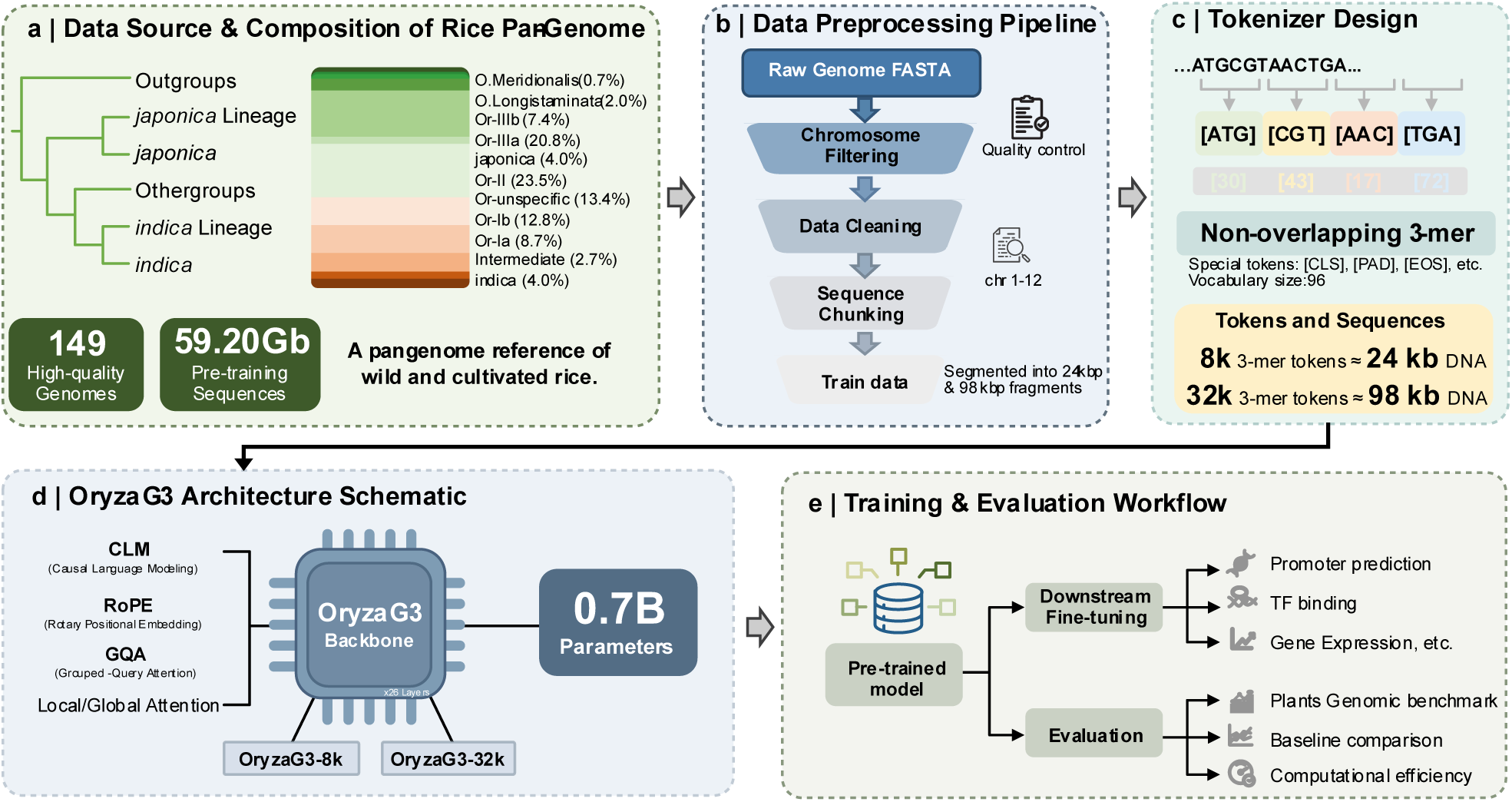
Overview of OryzaG3 model construction. (a) Rice pangenome data sources and composition. The pretraining corpus consists of 149 high-quality *Oryza* genomes, covering wild rice, cultivated rice, and outgroups, with a total sequence volume of 59.20 Gb. (b) Data preprocessing pipeline. Raw genomic data underwent chromosome filtering (retaining only chromosomes 1-12) and cleaning, then were segmented into approximately 24 kb and 98 kb fragments for training. (c) Tokenizer design. DNA sequences are segmented into non-overlapping 3-mer tokens, constructing a 96-token vocabulary containing 64 standard 3-mers and special reserved tokens. The figure shows that 8k and 32k tokens correspond to approximately 24 kb and 98 kb DNA sequence lengths, respectively. (d) Schematic of the OryzaG3 architecture. The model integrates causal language modeling, Rotary Position Embeddings (RoPE), and Grouped Query Attention (GQA), and provides 8k and 32k context-length versions. (e) Training and evaluation workflow. Pretrained models can be fine-tuned for downstream tasks such as promoter prediction, transcription factor (TF) binding, and gene expression; the model was also evaluated through the PGB, baseline comparisons, and computational efficiency analysis.

### OryzaG3 achieves competitive performance in PGB

To evaluate the downstream transferability of OryzaG3, we fine-tuned it on the rice polyA prediction task from the Plants Genomic Benchmark (including two subsets: oryza_sativa_indica_group and oryza_sativa_japonica_group) and compared it with AgroNT, Botanic0-L, and OneGenome-Rice baseline models. All models used unified training epochs, learning rates, optimizers, and evaluation metrics, with input token lengths set according to their respective tokenizers (see Materials and Methods for details).

In the oryza_sativa_indica_group task (see Fig. 2 and Table S1), OryzaG3-32k achieved an AUC of 0.970, AP of 0.941, F1 of 0.890, MCC of 0.835, and Accuracy of 0.926. OryzaG3-8k showed similar performance, with an AUC of 0.970, AP of 0.942, F1 of 0.887, and MCC of 0.830. Compared with multi-species models, OryzaG3-32k exhibited slightly higher AUC than AgroNT and Botanic0-L, along with modest MCC improvements. In the oryza_sativa_japonica_group task, OryzaG3 also achieved high predictive performance, although AgroNT was superior on some metrics (see Table S2). These results indicate that species-specific models pretrained on Oryza pangenome can attain performance comparable to or locally superior to larger multi-species models in rice-specific sequence classification tasks. Species-specific and multi-species models may offer complementary advantages: the former shows greater potential in efficiency and species adaptation, while the latter may gain broader generalization from cross-species corpora.

**Fig. 2.**
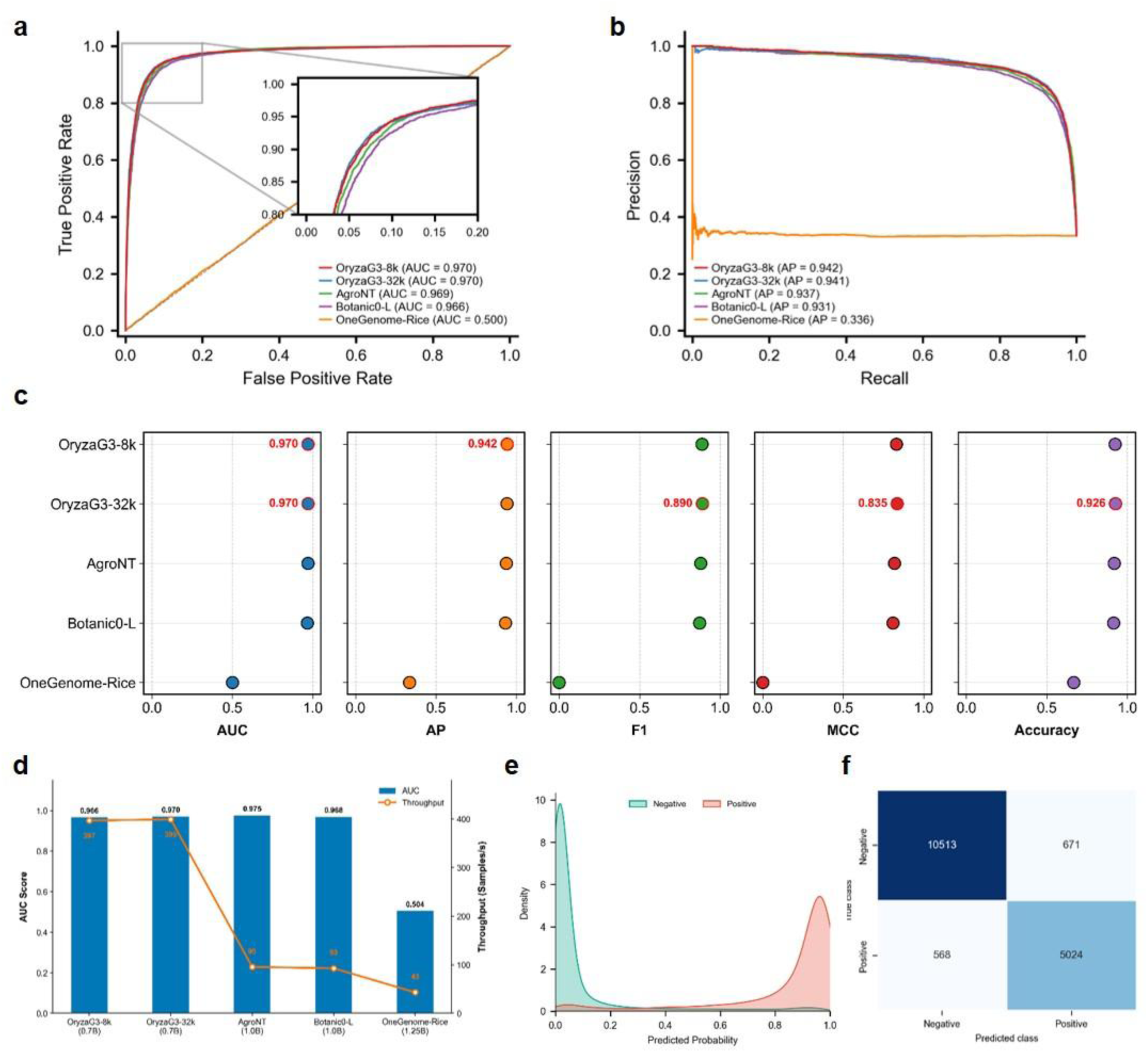
Performance and efficiency evaluation of OryzaG3 versus baseline models on the PGB polyA prediction task. (a) ROC curves of OryzaG3 (8k and 32k versions) and baseline models (AgroNT, Botanic0-L, OneGenome-Rice) on the test set of the polyA (oryza_sativa_indica_group) prediction task from the PGB. The inset shows details of the high true positive rate region; AUC values are provided in the legend. (b) Precision-Recall (PR) curves and Average Precision (AP) scores for the five models on the same test set. (c) Comprehensive comparison of the five models across AUC, AP, F1, MCC, and Accuracy. Red values highlight the best performance for each metric. (d) Comparison of model predictive performance and computational efficiency (test set prediction speed). Blue bars (left Y-axis) represent AUC scores; orange lines (right Y-axis) represent inference throughput (samples/s). The X-axis labels model names and parameter scales (in billions, B). (e) Prediction probability distribution of the OryzaG3-32k model. The kernel density estimation (KDE) plot shows the model’s discrimination between negative (green) and positive (red) samples. (f) Confusion matrix of OryzaG3-32k on the test set (classification threshold 0.5).

In terms of inference and computational efficiency, OryzaG3 also demonstrates clear advantages (see Fig. 2d, Table S1, and Table S2). Under identical testing conditions for the polyA prediction task (oryza_sativa_indica_group), inference throughput was evaluated on an NVIDIA A100-80GB GPU using the maximum batch size supported by each model. OryzaG3-32k achieved 399.80 samples/s, compared to 95.47 samples/s for AgroNT and 92.32 samples/s for Botanic0-L, corresponding to an approximately 4.2-4.3-fold improvement. This advantage likely stems from the smaller parameter scale, compact 3-mer vocabulary, and efficient model architecture. These results demonstrate that OryzaG3 can significantly reduce deployment costs for rice-specific tasks while maintaining competitive performance.

It should be noted that, because the input sequences in the PGB polyA dataset are relatively short, this benchmark cannot fully examine the advantages of the 32k context window. OryzaG3-32k is primarily designed for future long-context scenarios such as distal regulatory modeling, structural variant effect prediction, and chromosome-scale sequential representation learning. Dedicated long-sequence benchmarks are still needed to assess whether expanded context lengths yield biological predictive gains beyond short local sequence tasks. Additionally, under unified fine-tuning settings, the OneGenome-Rice model did not demonstrate effective classification performance on this task, consistent with the non-significant predictive results reported in its original study.¹⁴ This paper does not conclude that OneGenome-Rice lacks representational capacity; rather, it suggests that its MoE architecture, tokenization method, and pretraining objective may require more targeted fine-tuning strategies.

### Pretraining sufficiency significantly affects downstream task performance

To investigate the impact of pretraining progress on model representation quality, we extracted multiple checkpoints from early (E), middle (M), and late (L) stages during training (OryzaG3-8k-E/M/L and OryzaG3-32k-E/M/L; see Materials and Methods for details) and evaluated their performance on the polyA prediction task under identical fine-tuning parameters.

The results (Fig. 3, Table S3 and S4) show that early checkpoints have not yet formed effective transferable sequential representations. In the oryza_sativa_indica_group task, OryzaG3-8k-E achieved an AUC of only 0.586, with F1 and MCC close to 0, performing near random. In contrast, OryzaG3-8k-M showed significant improvement, with an AUC of 0.967, F1 of 0.882, and MCC of 0.821, approaching the fully pretrained OryzaG3-8k-L model. OryzaG3-8k-L further reached an AUC of 0.970, demonstrating more stable downstream transferability.

**Fig. 3.**
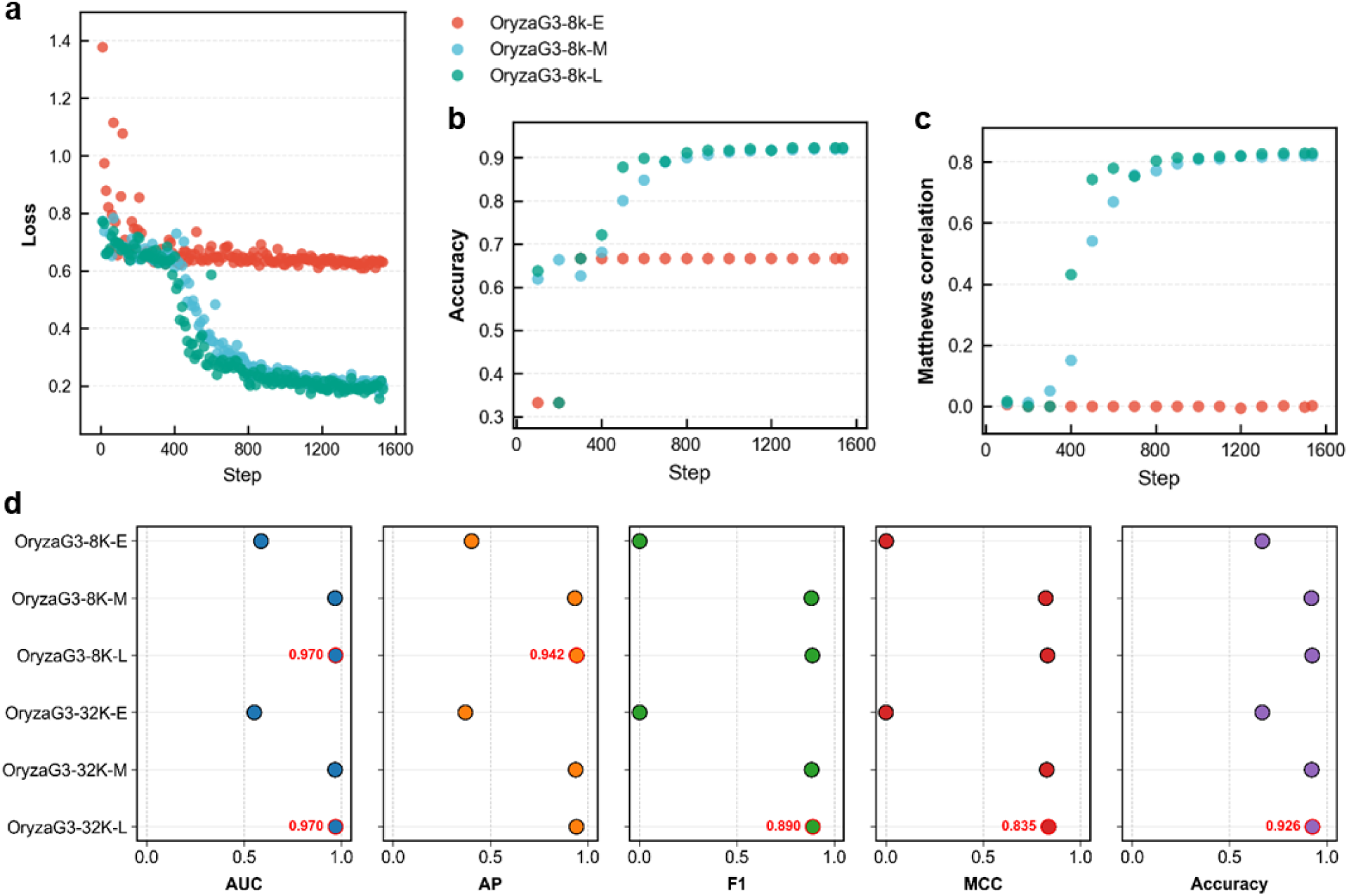
Impact of pretraining degree on OryzaG3 model performance. (a-c) Fine-tuning dynamics of OryzaG3-8k checkpoints from different pretraining stages (E: early, M: middle, L: late) on the polyA (oryza_sativa_indica_group) task, showing loss (a), accuracy (b), and MCC (c). Fully pretrained models (L stage) not only achieve superior performance but also converge faster (OryzaG3-32k results in Supplementary Materials). (d) Performance comparison of the two OryzaG3 configurations (8k, 32k) across three pretraining stages. Metrics include AUC, AP, F1, MCC, and Accuracy (test set: oryza_sativa_indica_group). Red markers and values highlight optimal performance.

These results indicate that models with only limited pretraining steps struggle to learn effective genomic sequential representations, whereas sufficient pretraining substantially enhances downstream performance. Importantly, fully pretrained models (stage L) converge faster during fine-tuning with more stable loss reduction (Fig. 3a-c), highlighting the critical importance of thorough traversal of high-quality pangenome data for species-specific model construction.

### Computational optimization for long-context training efficiency

Genomic sequence modeling typically faces the dual challenges of long-sequence inputs and high computational costs^6,7^. In particular, self-attention computation in long-context DNA modeling imposes substantial demands on memory and training time. Systematic comparisons of different optimization techniques remain limited in existing studies. To systematically evaluate the impact of various optimization techniques on training efficiency, we assessed four training strategies on NVIDIA A100-80GB GPUs across 8k, 16k, and 32k token context lengths: 1) Base: native Hugging Face Transformers^25^ implementation; 2) FA2: integrated FlashAttention-2^22^; 3) GC: enabled Gradient Checkpointing^23^; 4) FA2 + GC: combined use.

The results (Fig. 4) show that, for 8k context with batch size 1, the Base configuration required 2.50 s per step with 32.40 GB peak memory; enabling FA2 reduced time to 0.95 s (2.63× throughput improvement) and memory to 29.62 GB; GC increased time to 2.90 s but reduced memory to 9.41 GB; FA2 + GC achieved 1.24 s with 9.42 GB memory. At the more challenging 32k length with larger batch sizes, Base and single FA2 strategies readily triggered out-of-memory (OOM) errors, while FA2 + GC robustly controlled peak memory, supporting longer sequence modeling. The experiments demonstrate that FlashAttention-2 significantly accelerates training, while Gradient Checkpointing effectively reduces memory usage, with clear advantages in long-sequence scenarios. For short sequences (8k), FA2 is recommended; for long sequences (16k and above), the FA2 + GC combination achieves the optimal time-memory balance. The effects of different optimization strategies on training efficiency and memory usage are significant and complementary (detailed results in Table S5).

**Fig. 4.**
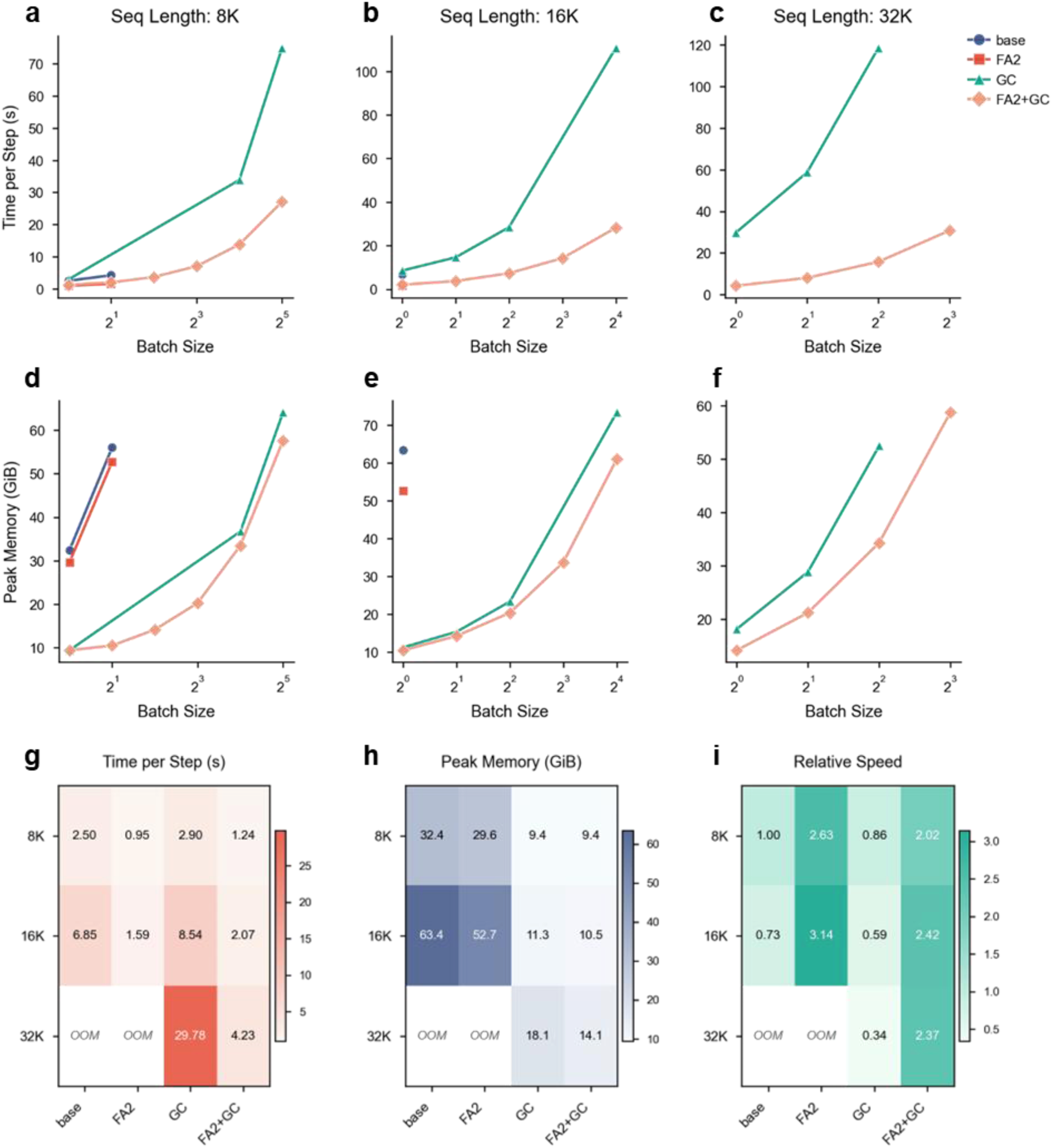
Comprehensive performance testing of memory-efficient training strategies. Systematic evaluation of Base, FA2 (FlashAttention-2), GC (Gradient Checkpointing), and FA2+GC across 8K, 16K, and 32K token sequence lengths for training time and memory consumption. (a-f) Scaling characteristics with batch size. (a-c) Training time per step (seconds) versus batch size for 8K, 16K, and 32K sequences. (d-f) Peak GPU memory consumption (GiB) under the same conditions. The X-axis uses a base-2 logarithmic scale for batch size. Points represent measured values; truncated or missing lines indicate training failures due to OOM. FA2+GC supports the largest batch sizes across all lengths. (g-i) Heatmaps of core metrics at batch size 1. (g) Time per step (s); (h) Peak memory (GiB); (i) Relative training speedup (normalized to Base at 8K = 1.00). Detailed experimental settings and results are in the Supplementary Materials.

Based on these systematic evaluations, we ultimately selected the 8k context version for primary pretraining and developed the 32k context version to support longer-sequence downstream tasks. This optimization strategy not only improved the development efficiency of OryzaG3 but also provides practical references for training genomic foundation models in other species.

## Discussion

In this study, we introduced OryzaG3, a single-species DNA language model trained on a comprehensive pangenome comprising 149 high-quality rice accessions. By utilizing a compact 3-mer vocabulary and a streamlined architecture of approximately 700 million parameters, OryzaG3 achieves competitive predictive accuracy on the rice polyA prediction task within the Plants Genomic Benchmark compared to leading multi-species models (AgroNT and Botanic0-L), while delivering substantial gains in inference throughput and computational velocity. These results establish a highly resource-efficient technical paradigm for building lightweight, species-specific genomic foundation models.

A key conceptual insight from OryzaG3 is that scaling parameter size is not the sole determinant of downstream performance in genomic language modeling. Instead, pretraining data quality, precise species-matching, optimized tokenization, and training sufficiency emerge as critical drivers of model behavior. By tracking the downstream efficacy of intermediate checkpoints, we empirically demonstrated that thorough traversal of the pangenomic data is essential; models evaluated at early training stages exhibited near-random performance, whereas fully pretrained variations unlocked superior accuracy and accelerated fine-tuning convergence. This suggests that future crop-specific foundational architectures should prioritize the comprehensive extraction of high-quality pangenomic variation and rigorous monitoring of training dynamics, rather than relying exclusively on brute-force parameter scaling.

From a computational perspective, long-context DNA modeling remains severely bottlenecked by hardware memory and training durations. Our efficiency ablation experiments reveal a clear functional division: FlashAttention-2 predominantly accelerates training velocity, whereas Gradient Checkpointing minimizes the memory footprint. Their synergy provides an optimal time-memory trade-off that enables long-context genomic training. Importantly, these algorithmic benchmarks extend beyond OryzaG3, offering immediate, actionable strategies for researchers training dedicated foundation models for other complex crop genomes, such as maize, wheat, and soybean.

By inheriting Rotary Position Embeddings (RoPE) and hybrid local-global attention mechanisms^18^, OryzaG3 is architecturally primed for ultra-long-context expansion. Given that key cis-regulatory interactions and structural variations in plant genomes often span vast genomic intervals, extending context lengths holds profound promise for distal element identification, structural variant characterization, and chromosome-scale sequence modeling^4,6–8^. However, because systematic validation on sequences exceeding 128k to 1M^+^ base pairs remains ongoing, these ultra-long applications should currently be interpreted as promising future directions rather than empirically consolidated conclusions of this work.

Despite these contributions, several limitations of the current framework warrant consideration. First, our systematic downstream validation focused primarily on the polyA prediction task; while this confirms OryzaG3’s robust sequence representation capabilities, it does not fully encompass the diverse landscape of plant functional genomics. Future work will expand this evaluation to broader agronomic tasks, including promoter and splice-site prediction, chromatin accessibility mapping, and variant effect assessment. Second, our interpretation of the coupling between training dynamics and underlying biological features remains in its preliminary stages, necessitating deeper integration with multi-omics layers. Finally, the incremental training protocols required for ultra-long contexts (>1 Mb) demand further algorithmic refinement and large-scale validation.

In conclusion, OryzaG3 validates a potent, “lightweight + species-specific” strategy fueled by high-quality pangenome resources. As an efficient computational engine, OryzaG3 is well-positioned to integrate with multi-omics annotations to rapidly empower downstream applications such as high-throughput variant effect screening and trait association mapping. As high-resolution pangenomic and functional genomic resources accumulate across diverse flora^26–28^, extending this framework to other crops will accelerate the paradigm shift from baseline sequence understanding to targeted functional prediction and predictive design breeding.

## Materials and Methods

### Rice pangenome dataset and preprocessing

The pretraining corpus for this study was derived from publicly available high-quality rice pangenome resources (NGDC BioProject accession: PRJCA024131; genome details in Table S6). The dataset comprises 149 high-quality *Oryza* pangenome, broadly covering cultivated rice (*Oryza sativa*), wild rice (*Oryza rufipogon*), and related outgroup species, with a total raw data volume of 59.20 Gb.

This study extracted only sequences from chromosomes 1-12. For the small number of unknown bases (N) present in high-quality assemblies, a random substitution strategy was applied, replacing each N with one of A, T, G, or C. During sequence segmentation, chromosome boundaries were strictly respected. Long sequences were segmented using sliding windows of fixed lengths 24,570 bp and 98,298 bp, with terminal fragments shorter than the fixed length retained. To improve subsequent data loading and tokenization efficiency, a pre-tokenization step was performed during data preprocessing by inserting a space every three bases. Ultimately, the 24,570 bp corpus for pretraining contained 2.42 million sequences, and the 98,298 bp corpus contained 606.65K sequences (see Table S7 for details).

### 3-mer tokenizer construction

OryzaG3 employs a non-overlapping 3-mer tokenization strategy and constructs a DNA-specific tokenizer (ATGC-3mer-withUnused_tokenizer) from scratch. The tokenizer vocabulary is compact, with a size of 96, including 64 standard 3-mer combinations and necessary special reserved tokens. For input DNA sequences, the model segments them into triplet tokens with a 3-bp stride.

The choice of a non-overlapping 3-mer tokenizer was based on the following considerations. First, the small 3-mer vocabulary significantly reduces the number of embedding parameters and associated computational overhead. Second, the triplet scale offers biological interpretability, corresponding to codon length. Third, non-overlapping segmentation preserves local sequence information while reducing input token length, thereby improving modeling efficiency for long sequences. The tokenizer construction code is available in the Code and data availability section.

### OryzaG3 model architecture

OryzaG3 is built upon the Gemma 3-1B architecture (config), with the original large natural language vocabulary replaced by the DNA-specific 3-mer vocabulary described above, substantially reducing embedding layer parameters. The model backbone consists of 26 Transformer blocks, with a hidden size of 1,152, 4 attention heads, and a head dimension of 256. The model integrates Rotary Position Embeddings (RoPE) and retains Gemma3 hybrid local-global attention mechanism, resulting in a total parameter count of approximately 700M. Detailed model configurations are provided in Table S8.

Two context-length versions were constructed: OryzaG3-8k and OryzaG3-32k, with maximum context lengths of 8,192 tokens and 32,768 tokens, respectively. Both versions use the same pretraining corpus, the same 3-mer tokenizer, and the same backbone architecture. The primary differences lie in maximum context length, corresponding training sample segmentation lengths, and batch size configurations. Detailed model configurations are available in the Code and data availability section.

### Pretraining objective and optimization

OryzaG3 adopts CLM as the self-supervised pretraining objective. Given an input sequence of 3-mer tokens, the model predicts the next token based on all preceding tokens, thereby learning local combinatorial patterns, contextual dependencies, and potential functional sequence features in DNA. Pretraining utilized the AdamW optimizer with an initial learning rate of 2 × 10⁻⁴ and a cosine decay learning rate schedule. Training was performed with bfloat16 mixed precision to reduce memory usage and improve efficiency. Pretraining was completed on a server equipped with 8 NVIDIA H100-80GB GPUs. Key training hyperparameters, batch sizes, training steps, and resource configurations are detailed in Table S9 and the Code and data availability section.

OryzaG3-8k and OryzaG3-32k were pretrained using data corresponding to their respective context lengths. In terms of sequence length conversion: for 8k context pretraining, the original 24,570 bp DNA sequences were converted into 8,190 3-mer tokens; for 32k context pretraining, the original 98,298 bp sequences were converted into 32,766 3-mer tokens. Remaining token positions were filled with special padding tokens.

### Plants Genomic Benchmark fine-tuning tasks

Downstream task evaluation was performed using the rice polyA prediction task from the PGB. Two rice-related subsets were used: poly_a.oryza_sativa_indica_group and poly_a.oryza_sativa_japonica_group. PGB raw data were provided in FASTA format. For compatibility with a unified fine-tuning pipeline, the data were converted into CSV files containing sequence and label columns. Labels are binary, indicating whether the input sequence corresponds to a polyA -related positive sample. The datasets and conversion scripts are available in the Code and data availability section.

The oryza_sativa_indica_group task contains 98,139 training samples and 16,776 test samples; the oryza_sativa_japonica_group task contains 120,621 training samples and 20,232 test samples. Each original input sequence is 400 bp in length. Both training and test sets follow the official PGB splits without additional re-partitioning. Detailed data statistics are provided in Table S10.

### Baseline models and fine-tuning settings

AgroNT, Botanic0-L, and OneGenome-Rice were selected as baseline models for comparison. All models were loaded with their official open-source pretrained weights, tokenized using their original tokenizers, and equipped with a newly initialized mean-pooling classification head. To ensure fair comparison, all models used the same random seed, official data splits, training epochs (2 epochs), optimizer (AdamW), learning rate, and weight decay strategy during fine-tuning. Batch size was adaptively set to the maximum feasible capacity within each model’s memory constraints (see Table S11).

Owing to differences in tokenizer granularity across models, the actual token length for fixed-length input DNA sequences was calculated and rounded up to the nearest multiple of 8 to accommodate Transformer hardware acceleration. The input token length was computed as follows:

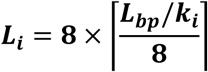

where *L*_*i*_ represents the input token length for the *i*-th model, *L*_bp_ is the input DNA sequence length (set to 512 bp in the polyA task to cover the 400 bp sequence plus special tokens), and *k*_*i*_ is the granularity of the corresponding tokenizer. According to this calculation, a 512 bp input corresponds to 176 tokens for OryzaG3, 88 tokens for AgroNT and Botanic0-L, and 512 tokens for OneGenome-Rice.

For computational efficiency evaluation, inference speed (throughput) was measured based on inference time on the test set. Since no weight updates occur during testing, this metric accurately reflects the inference efficiency of each model in real-world deployment scenarios.

### Checkpoint selection and pretraining diagnostics

To systematically evaluate the impact of pretraining progress on the model’s transferable representation learning, multiple checkpoints were saved during pretraining and categorized into early, middle, and late stages according to training progress. The corresponding models were named OryzaG3-8k-E, OryzaG3-8k-M, OryzaG3-8k-L, OryzaG3-32k-E, OryzaG3-32k-M, and OryzaG3-32k-L.

All checkpoints were evaluated under identical downstream fine-tuning settings to compare their transferable representational capabilities. The evaluation task was the rice polyA prediction task from the Plants Genomic Benchmark. Specific checkpoint selection points are detailed in Table S12.

### Computational optimization for long-context training efficiency

Computational efficiency ablation experiments were conducted on NVIDIA A100-80GB GPUs. Four training strategies were systematically compared: (1) Base: native Hugging Face Transformers implementation; (2) FlashAttention-2 (FA2): FlashAttention-2 enabled on top of Hugging Face Transformers; (3) Gradient Checkpointing (GC): Gradient Checkpointing enabled on top of Hugging Face Transformers; (4) FA2 + GC: both FlashAttention-2 and Gradient Checkpointing enabled simultaneously. Experiments were performed at three context lengths: 8,192 (8k), 16,384 (16k), and 32,768 (32k) tokens. For each configuration, batch size was incrementally increased in powers of 2 while staying within memory limits to systematically assess the impact of different strategies on computational efficiency and memory usage.

Training time (Time per step) was recorded from the step time output by the training code. Peak memory usage was obtained from the maximum GPU memory value reported by nvidia-smi during execution (in MiB). Relative training speed was calculated with the Base strategy at 8k context and batch size = 1 normalized to 1.00. When out-of-memory errors prevented training completion, the corresponding entry was marked as “-”. Complete results are provided in Table S5.

### Hardware and software environment

All model pretraining was conducted on a computing cluster equipped with NVIDIA H100-80GB GPUs. Downstream task fine-tuning evaluations and long-context training efficiency ablation experiments were performed on hardware equipped with NVIDIA A100-80GB GPUs.

### Evaluation metrics

For binary classification tasks, model performance was evaluated using AUC, Average Precision (AP), Accuracy, F1-score, Precision, Recall, and Matthews Correlation Coefficient (MCC). AUC and AP were computed based on the model’s predicted probabilities. Accuracy, F1-score, Precision, Recall, and MCC were calculated based on binary predicted labels. Unless otherwise specified, the default classification threshold was set to 0.5. Descriptions of each evaluation metric are provided in Table S13.

## Code and data availability

Model weights: The pretrained OryzaG3 models are available on Hugging Face.

https://huggingface.co/yang0104/OryzaG3-8k

https://huggingface.co/yang0104/OryzaG3-32k

Code repository: The complete codebase, including data preprocessing, tokenizer scripts, pretraining and fine-tuning configurations, is available on GitHub.

(link to be provided upon formal publication)

Pretraining data: The 149 rice pangenome are from a 2025 Nature paper and are deposited in the National Genomics Data Center (NGDC), BioProject PRJCA024131.

https://ngdc.cncb.ac.cn/bioproject/browse/PRJCA024131

Fine-tuning benchmark data: The polyA prediction task data from the Plants Genomic Benchmark were downloaded from the Hugging Face repository InstaDeepAI/plant-genomic-benchmark.

https://huggingface.co/datasets/InstaDeepAI/plant-genomic-benchmark

Gemma 3 architecture configuration: Obtained from “google/gemma-3-1b-pt” on Hugging Face. https://huggingface.co/google/gemma-3-1b-pt

Baseline model access paths:

AgroNT: Obtained from “InstaDeepAI/agro-nucleotide-transformer-1b” on Hugging Face.

https://huggingface.co/InstaDeepAI/agro-nucleotide-transformer-1b

Botanic0: Obtained from “living-models/Botanic0-L” on Hugging Face.

https://huggingface.co/living-models/Botanic0-L

OneGenome-Rice: Obtained from “ZhejiangLab/OneGenome-Rice” on Hugging Face.

https://huggingface.co/ZhejiangLab/OneGenome-Rice

## Author contributions

Z.Xia conceived the study. L.Yang, Y.Xia, Z.Yang, C.Xia, T.Wu and M.Zou, contributed to the design, training, and programming of the OryzaG3 model. L.Yang wrote the manuscript with input from all authors. All authors read and approved the final manuscript.

## Acknowledgements

We also acknowledge the developers of Gemma, FlashAttention, HuggingFace Transformers and the Plants Genomic Benchmark for making their resources publicly available.

## Funding Statements

This work was supported by Biological Breeding-National Science and Technology Major Project (2023ZD04073), Guangxi Ministry of Science and Technology (GuikeAA23062015), and High-performance Computing Platform of YZBSTCACC.

## Supplementary Materials

### Supplementary Tables

**Table S1.** Fine-tuning performance comparison of OryzaG3 and baseline models on the polyA prediction task (oryza_sativa_indica_group)

| Model | AUC | AP | F1 | MCC | Accuracy | Runtime (s) | Samples/s |
| --- | --- | --- | --- | --- | --- | --- | --- |
| OryzaG3-8k (700M) | 0.970 | 0.942 | 0.887 | 0.830 | 0.924 | 41.90 | 400.41 |
| OryzaG3-32k (700M) | 0.970 | 0.941 | 0.890 | 0.835 | 0.926 | 41.96 | 399.80 |
| AgroNT (1B) | 0.969 | 0.937 | 0.879 | 0.818 | 0.919 | 175.71 | 95.47 |
| Botanic0-L (991M) | 0.966 | 0.931 | 0.873 | 0.809 | 0.914 | 181.72 | 92.32 |
| OneGenome-Rice(1.25B) | 0.500 | 0.336 | - | - | 0.667 | 391.30 | 42.87 |

Note: Runtime excludes tokenization and data loading time; F1 and MCC were calculated using a 0.5 threshold; “-” indicates that the metric could not be computed or equals zero.

**Table S2.** Fine-tuning performance comparison of OryzaG3 and baseline models on the polyA prediction task (oryza_sativa_japonica_group)

| Model | AUC | AP | F1 | MCC | Accuracy | Runtime (s) | Samples/s |
| --- | --- | --- | --- | --- | --- | --- | --- |
| OryzaG3-8k (700M) | 0.966 | 0.936 | 0.882 | 0.823 | 0.921 | 51.00 | 396.72 |
| OryzaG3-32k (700M) | 0.970 | 0.945 | 0.887 | 0.830 | 0.924 | 50.67 | 399.26 |
| AgroNT (1B) | 0.975 | 0.952 | 0.900 | 0.849 | 0.933 | 212.02 | 95.42 |
| Botanic0-L (991M) | 0.968 | 0.940 | 0.884 | 0.825 | 0.921 | 218.49 | 92.60 |
| OneGenome-Rice(1.25B) | 0.504 | 0.336 | - | - | 0.667 | 470.32 | 43.02 |

Note: Runtime excludes tokenization and data loading time; F1 and MCC were calculated using a 0.5 threshold; “-” indicates that the metric could not be computed or equals zero.

**Table S3.** Fine-tuning performance comparison of OryzaG3 models at different pretraining stages on the polyA prediction task (oryza_sativa_indica_group)

| Models | AUC | AP | F1 | MCC | Accuracy |
| --- | --- | --- | --- | --- | --- |
| OryzaG3-8k-E | 0.586 | 0.403 | 0.001 | 0.002 | 0.667 |
| OryzaG3-8k-M | 0.967 | 0.933 | 0.882 | 0.821 | 0.920 |
| OryzaG3-8k-L | 0.970 | 0.942 | 0.887 | 0.830 | 0.924 |
| OryzaG3-32k-E | 0.552 | 0.371 | - | - | 0.667 |
| OryzaG3-32k-M | 0.967 | 0.937 | 0.884 | 0.826 | 0.922 |
| OryzaG3-32k-L | 0.970 | 0.941 | 0.890 | 0.835 | 0.926 |

Note: “-” indicates that the metric could not be computed or equals zero.

**Table S4.** Fine-tuning performance comparison of OryzaG3 models at different pretraining stages on the polyA prediction task (oryza_sativa_japonica_group)

| Models | AUC | AP | F1 | MCC | Accuracy |
| --- | --- | --- | --- | --- | --- |
| OryzaG3-8k-E | 0.890 | 0.780 | 0.764 | 0.637 | 0.830 |
| OryzaG3-8k-M | 0.971 | 0.947 | 0.890 | 0.835 | 0.926 |
| OryzaG3-8k-L | 0.966 | 0.936 | 0.882 | 0.823 | 0.921 |
| OryzaG3-32k-E | 0.831 | 0.669 | 0.553 | 0.408 | 0.753 |
| OryzaG3-32k-M | 0.970 | 0.946 | 0.890 | 0.834 | 0.926 |
| OryzaG3-32k-L | 0.970 | 0.945 | 0.887 | 0.830 | 0.924 |

**Table S5.** Training efficiency under different memory-optimization configurations.

| Treatment | token_length | batch_size | Time per steps (s) | Peak memory (Mib) | Relative speed |
| --- | --- | --- | --- | --- | --- |
| base | 8192 | 1 | 2.50 | 33183 | 1.00 |
| base | 8192 | 2 | 4.21 | 57381 | 1.19 |
| base | 8192 | 4 | - | OOM | - |
| base | 16384 | 1 | 6.85 | 64919 | 0.73 |
| base | 32786 | 1 | - | OOM | - |
| FA2 | 8192 | 1 | 0.95 | 30331 | 2.63 |
| FA2 | 8192 | 2 | 1.55 | 54035 | 3.23 |
| FA2 | 8192 | 4 | - | OOM | - |
| FA2 | 16384 | 1 | 1.59 | 53939 | 3.14 |
| FA2 | 16384 | 2 | - | OOM | - |
| FA2 | 32786 | 1 | - | OOM | - |
| GC | 8192 | 1 | 2.90 | 9635 | 0.86 |
| GC | 8192 | 16 | 33.88 | 37565 | 1.18 |
| GC | 8192 | 32 | 74.97 | 65589 | 1.07 |
| GC | 8192 | 64 | - | OOM | - |
| GC | 16384 | 1 | 8.54 | 11521 | 0.59 |
| GC | 16384 | 2 | 14.62 | 15753 | 0.68 |
| GC | 16384 | 4 | 28.43 | 23937 | 0.70 |
| GC | 16384 | 16 | 110.74 | 75125 | 0.72 |
| GC | 16384 | 32 | - | OOM | - |
| GC | 32786 | 1 | 29.78 | 18553 | 0.34 |
| GC | 32786 | 2 | 58.76 | 29521 | 0.34 |
| GC | 32786 | 4 | 118.60 | 53725 | 0.34 |
| GC | 32786 | 8 | - | OOM | - |
| FA2+GC | 8192 | 1 | 1.24 | 9643 | 2.02 |
| FA2+GC | 8192 | 2 | 1.97 | 10757 | 2.54 |
| FA2+GC | 8192 | 4 | 3.68 | 14497 | 2.72 |
| FA2+GC | 8192 | 8 | 7.03 | 20753 | 2.84 |
| FA2+GC | 8192 | 16 | 13.78 | 34269 | 2.90 |
| FA2+GC | 8192 | 32 | 27.11 | 58997 | 2.95 |
| FA2+GC | 8192 | 64 | - | OOM | - |
| FA2+GC | 16384 | 1 | 2.07 | 10749 | 2.42 |
| FA2+GC | 16384 | 2 | 3.76 | 14617 | 2.66 |
| FA2+GC | 16384 | 4 | 7.31 | 20897 | 2.74 |
| FA2+GC | 16384 | 8 | 14.26 | 34557 | 2.81 |
| FA2+GC | 16384 | 16 | 28.21 | 62453 | 2.84 |
| FA2+GC | 16384 | 32 | - | OOM | - |
| FA2+GC | 32786 | 1 | 4.23 | 14473 | 2.37 |
| FA2+GC | 32786 | 2 | 8.02 | 21681 | 2.50 |
| FA2+GC | 32786 | 4 | 15.75 | 35133 | 2.54 |
| FA2+GC | 32786 | 8 | 30.79 | 60149 | 2.60 |
| FA2+GC | 32786 | 16 | - | OOM | - |

Note: OOM indicates out-of-memory error, preventing computation; “-” indicates that the value could not be computed due to OOM.

**Table S6.**
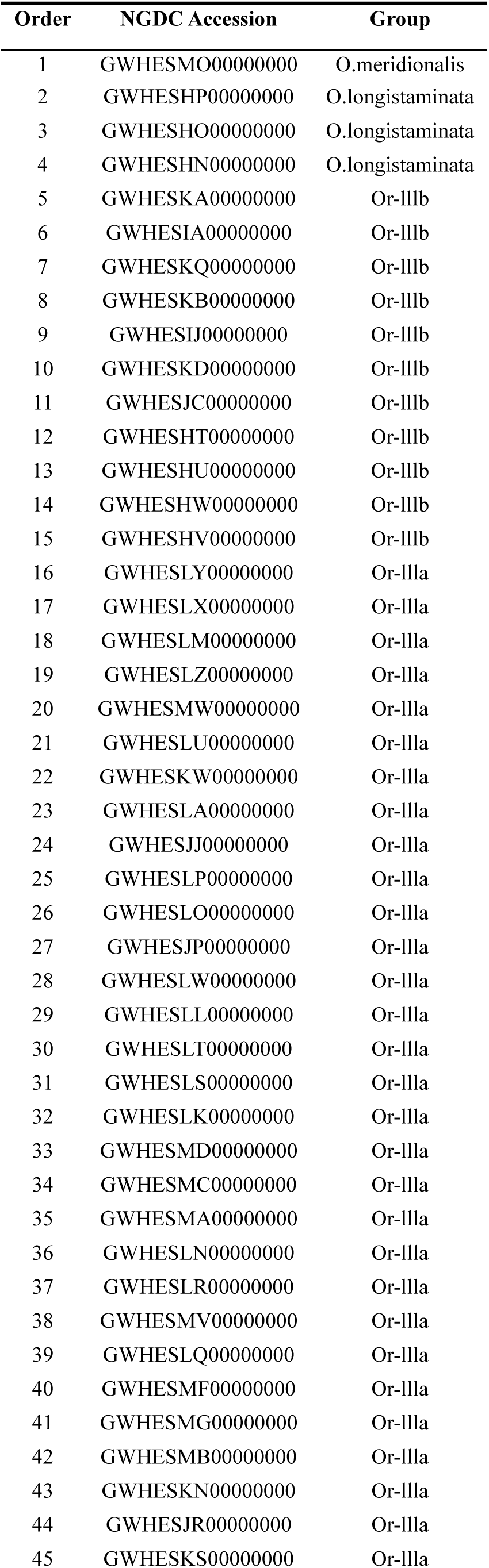
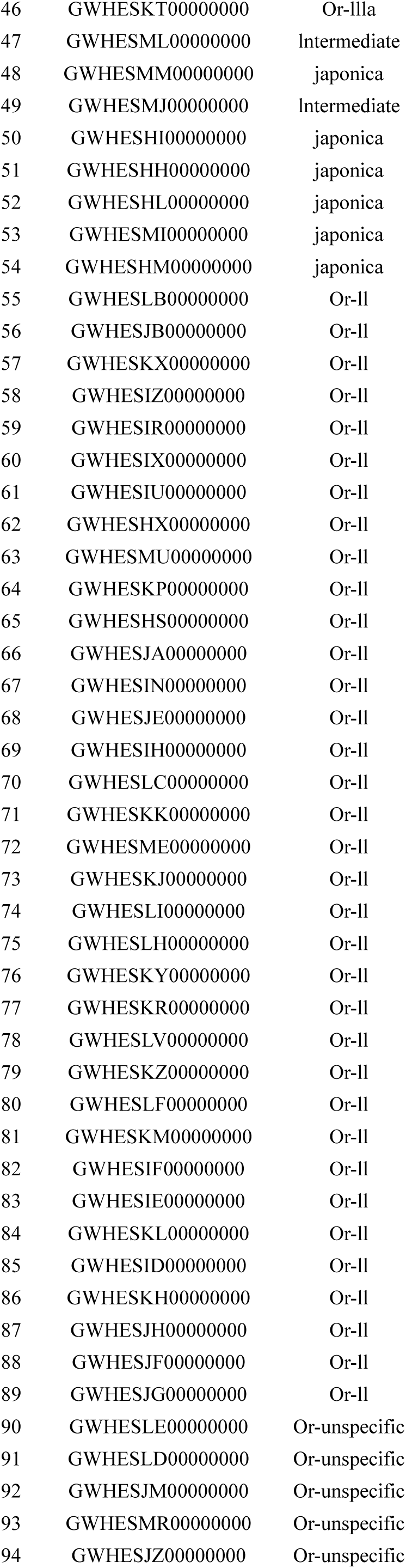
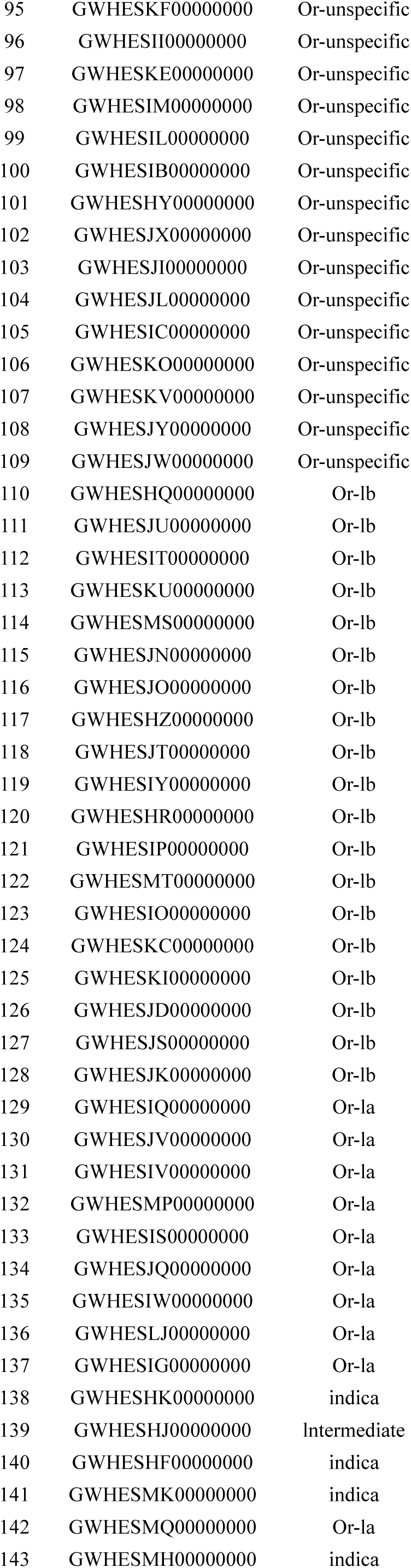
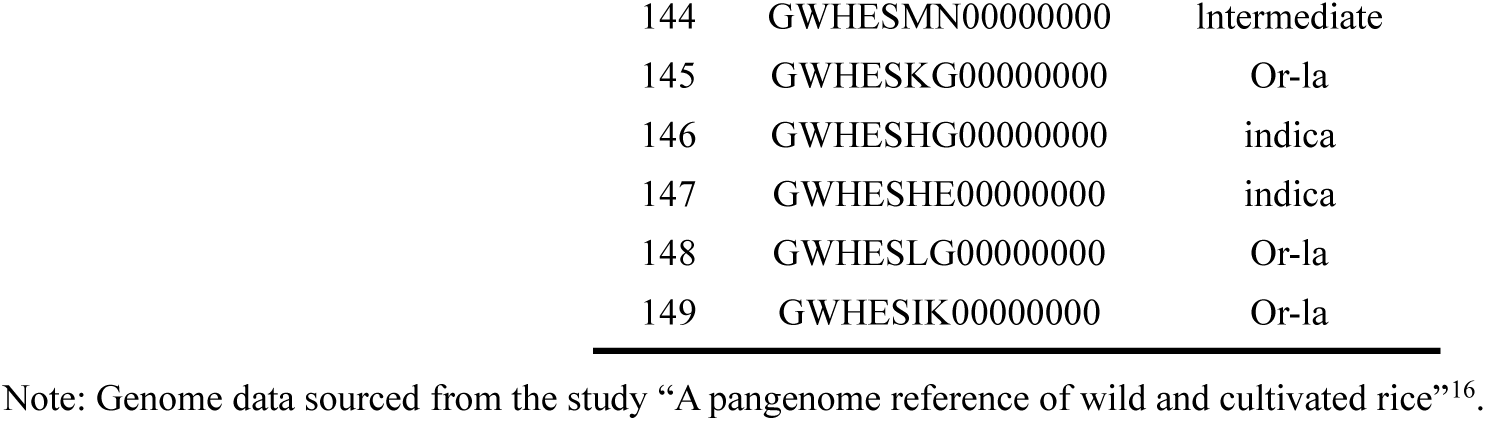
Information on the 149 rice pangenome and their sources.

**Table S7.** Characteristics of the pretraining data.

| Parameter | Value |
| --- | --- |
| Species | Rice(*Oryza*) |
| Number of Genomes | 149 |
| Number of Rice Groups | 11 |
| Total Data Size | 59.20GB |
| Genome Coverage | Chromosomes 1-12 |
| Training Samples (segmented at 24K bp) | 2.42 M |
| Training Samples (segmented at 98K bp) | 606.65 K |

**Table S8.** Main hyperparameters of the OryzaG3 model architecture.

| Parameter | Value |
| --- | --- |
| Hidden Size | 1152 |
| Num Hidden Layers | 26 |
| Num Attention Heads | 4 |
| Num Key-Value Heads | 1 |
| Intermediate Size | 6912 |
| Head Dim | 256 |
| Vocab Size | 96 |
| Max Position Embeddings | 32768 |
| Position Encoding | RoPE |
| Rope Theta | 1,000,000 |
| Attention Mechanism | Hybrid |
| Sliding Window Pattern | 6 |
| Sliding Window Size | 512 |
| Activation Function | gelu_pytorch_tanh |
| Torch Dtype | bfloat16 |

Note: The OryzaG3 model was initialized using only the Gemma3-1B architecture (config), without loading the original pretrained weights. The embedding layer was reinitialized with a DNA-specific 3-mer vocabulary (see the open-source model files on Hugging Face for implementation details).

**Table S9.** Main hyperparameters for OryzaG3 pretraining.

| Parameter | OryzaG3-8k | OryzaG3-32k |
| --- | --- | --- |
| Training Framework | Hugging Face-Transformers | Hugging Face-Transformers |
| Attention Implementation | FlashAttention-2 | FlashAttention-2 |
| Gradient Checkpointing | True | True |
| Context Length (tokens) | 8192 | 32768 |
| Per Device Batch Size | 32 | 8 |
| Gradient Accumulation Steps | 2 | 2 |
| Num Train Epochs | 1 | 1 |
| Learning Rate | $2 \times 10^{-4}$ | $2 \times 10^{-4}$ |
| LR Scheduler | Cosine | Cosine |
| Mixed Precision | bfloat16 | bfloat16 |
| Optimizer | AdamW | AdamW |

**Table S10.**
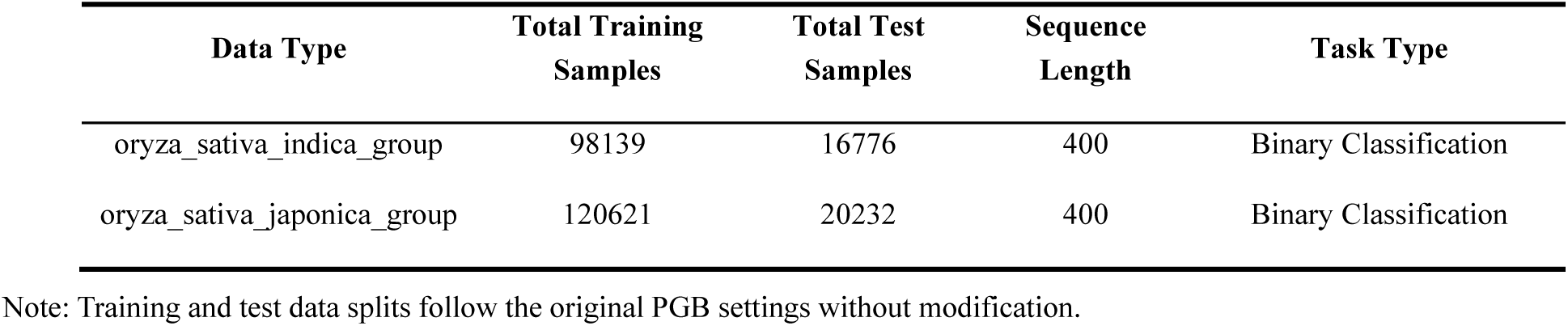
Characteristics of the fine-tuning datasets.

| Data Type | Total Training Samples | Total Test Samples | Sequence Length | Task Type |
| --- | --- | --- | --- | --- |
| oryza_sativa_indica_group | 98139 | 16776 | 400 | Binary Classification |
| oryza_sativa_japonica_group | 120621 | 20232 | 400 | Binary Classification |

Note: Training and test data splits follow the original PGB settings without modification.

**Table S11.** Main hyperparameters for OryzaG3 fine-tuning on the polyA prediction task.

| Base Model | OryzaG3 (700M) | AgroNT (1B) | Botanic0-L (991M) | OneGenome-Rice (1.25B) |
| --- | --- | --- | --- | --- |
| Tokenization | 3-mer | 3-mer | 6-mer | Singe-base |
| Context Length | 176 | 88 | 88 | 512 |
| Re-init Classifier Layer | True | True | True | True |
| Per Device Train Batch Size | 128 | 124 | 124 | 36 |
| Per Device Eval Batch Size | 128 | 124 | 124 | 36 |
| Gradient Accumulation Steps | 1 | 1 | 1 | 1 |
| Num Train Epochs | 2 | 2 | 2 | 2 |
| Learning Rate | 1e-4 | 1e-4 | 1e-4 | 1e-4 |
| Warmup Ratio | 0.05 | 0.05 | 0.05 | 0.05 |
| Weight Decay | 0.01 | 0.01 | 0.01 | 0.01 |
| Optimizer | AdamW | AdamW | AdamW | AdamW |

Note: Context Length was calculated according to tokenizer granularity; Batch Size was adjusted based on GPU memory capacity
(affected by differences in token length and model parameter size). All other hyperparameters were kept identical.

**Table S12.** Correspondence table of OryzaG3 pretraining checkpoints.

| Models | Checkpoint |
| --- | --- |
| OryzaG3-8k-E | checkpoint-30 |
| OryzaG3-8k-M | checkpoint-2010 |
| OryzaG3-8k-L | checkpoint-4735 |
| OryzaG3-32k-E | checkpoint-30 |
| OryzaG3-32k-M | checkpoint-2010 |
| OryzaG3-32k-L | checkpoint-4740 |

Note: The total number of training steps for OryzaG3-8k was 4,735, and for OryzaG3-32k was 4,740.

**Table S13.** Description of evaluation metrics.

| Metric | Description |
| --- | --- |
| Loss | The loss function measures the discrepancy between the model's predictions and the true labels. A smaller value indicates a stronger ability of the model to fit the test data and a lower prediction error. |
| Accuracy | The proportion of correctly classified samples out of the total test samples. Formula:<br>$\frac{TP+TN}{TP+TN+FP+FN}$ |
| F1 | The F1 score is the harmonic mean of Precision and Recall. Formula: $2 \times \frac{\text{Precision} \times \text{Recall}}{\text{Precision} + \text{Recall}}$ |
| Precision | The proportion of samples predicted as positive that are truly positive. Formula: $\frac{TP}{TP+FP}$ |
| Recall | The proportion of actually positive samples that are correctly predicted as positive. Formula:<br>$\frac{TP}{TP+FN}$ |
| Matthews Correlation Coefficient<br>(MCC) | A balanced binary classification metric that takes all four categories of the confusion matrix (TP, TN, FP, FN) into account. It ranges from -1 to 1, where 1 indicates a perfect prediction, 0 indicates a random guess, and -1 indicates complete disagreement. MCC provides a relatively objective evaluation even under extreme class imbalance. Formula: $\text{MCC} = \frac{TP \times TN - FP \times FN}{\sqrt{(TP+FP)(TP+FN)(TN+FP)(TN+FN)}}$ |
| Area Under the ROC Curve<br>(AUC / ROC-AUC) | The area under the ROC curve, which plots the False Positive Rate (FPR) on the x-axis and the True Positive Rate (TPR, i.e., Recall) on the y-axis. AUC measures the model's ability to distinguish between positive and negative classes. It ranges from 0.5 to 1, where 0.5 indicates no discriminative ability (random classification) and 1 indicates perfect discrimination. AUC is insensitive to class imbalance and is an important metric for binary classification. |
| Average Precision<br>(AP) | The average precision, i.e., the area under the Precision-Recall curve. It is typically computed as a weighted average of precision values at different recall thresholds. |

## Notes

### Competing Interest Statement

The authors have declared no competing interest.

